# Automated quantification of synaptic boutons reveals their 3D distribution in the honey bee mushroom body

**DOI:** 10.1101/589770

**Authors:** Amélie Cabirol, Albrecht Haase

## Abstract

Synaptic boutons are highly plastic structures undergoing experience-dependent changes in their number, volume, and shape. Their plasticity has been intensively studied in the insect mushroom bodies by manually counting the number of boutons in small regions of interest and extrapolating this number to the volume of the mushroom body neuropil.

Here we extend this analysis to the synaptic bouton distribution within a larger subregion of the mushroom body olfactory neuropil of honey bees (*Apis mellifera*). This required the development of an automated method combining two-photon imaging with advanced image post-processing and multiple threshold segmentation. The method was first validated in subregions of the mushroom body olfactory and visual neuropils. Further analyses in the olfactory neuropil suggested that previous studies overestimated the number of synaptic boutons. As a reason for that, we identified boundaries effects in the small volume samples. The application of the automated analysis to larger volumes of the mushroom body olfactory neuropil revealed a corrected average density of synaptic boutons and, for the first time, their 3D spatial distribution. This distribution exhibited a considerable heterogeneity.

This additional information on the synaptic bouton distribution provides the basis for future studies on brain development, symmetry, and plasticity.

## Introduction

Since the French biologist Félix Dujardin first described the insect mushroom bodies (MBs)^1^, these paired brain structures have been intensively studied due to their involvement in multisensory processing, learning, and memory^2–5^. The structural and functional plasticity occurring at their input region, the calyces, has been of particular interest in the past decade thanks to technological advances in fluorescence microscopy^6–9^.

Within the MB calyces, synaptic complexes, called microglomeruli, are formed by the terminals of neurons coming from the first sensory brain centers and the dendrites of MB neurons^4,10,11^. In honey bees and other Hymenoptera, information from different sensory modalities is segregated between three subregions of the MB calyx: the lip receives olfactory afferents from the antennal lobes, the collar receives visual afferents from the optic lobes, and the basal ring receives both olfactory and visual afferents^4^. The number of microglomeruli in these MB subregions has been shown to vary in response to sensory stimulations^12–15^, to memorization of olfactory and visual information^13,16,17^, and to aging^8,18^. Such variations have consequences on cognitive performance^19–21^.

In most studies, counting of the microglomeruli was performed manually in small regions of interest (ROIs) after immunostaining synapsin, a protein located in the axon terminals of neurons^8^. Manual counting is an adequate method to study relative changes in the microglomerular density. However, it is not only time-consuming but also subjective and therefore requires a blinded analysis. It is also prone to errors since it assumes that microglomeruli are homogeneously distributed within the MB calyx by using small ROIs to estimate an overall average density^22^. Since there is no efficient automated analysis of larger ROIs, information on the spatial distribution of microglomeruli within the MB calyx has been missing.

A few automated methods have been suggested but were either applied to small ROIs only^23^, or their performance has been seriously questioned^24,25^ because the absolute numbers of microglomeruli they delivered deviated strongly from those previously reported. An obvious limitation of automated counting is the proximity of microglomeruli within the MBs, which is at the resolution limit of those microscope objectives that would allow imaging larger regions of the MBs. Moreover, the heterogeneity of fluorescent dye distribution after immunohistochemical staining of whole-mounted brains prevents simple standard segmentation algorithms to obtain accurate counting results.

The objective of this study was to quantify the numbers of microglomeruli in larger regions of interest in the MB lip by developing a new automated analysis. The automated procedure reduced subjectivity and accelerated the evaluation considerably. Beyond that, the technique provided completely novel information on the 3D distribution of microglomeruli in a subregion of the MB olfactory neuropil. This will allow the search for stereotypical patterns in microglomerular density and the investigation of the heterogeneity of non-associative and associative plasticity induced by different learning paradigms and sensory exposure.

## Results and Discussion

The validity of the automated method was first assessed in two subregions of the MB calyx: the olfactory lip and the visual dense collar (Fig. 1A). The method was then applied to a 40-μm-thick subregion of the MB lip (Fig. 1B) to investigate the relationship between the number of microglomeruli and the ROI volume and thus estimate the number of microglomeruli in the whole lip. Finally, the 3D distribution of microglomeruli in the 40-μm-thick subregion of the MB lip was characterized.

**Figure 1.**
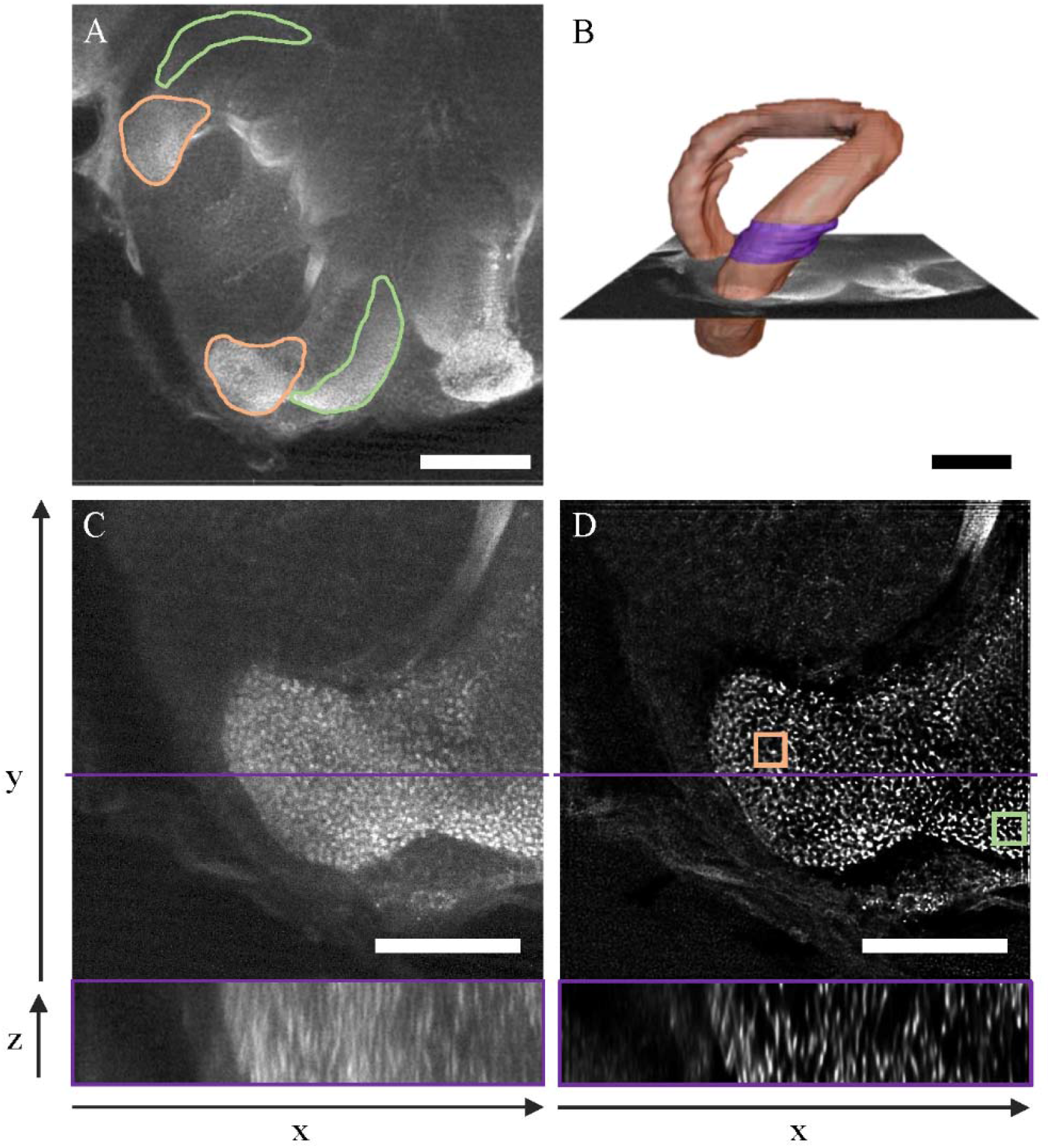
Post-acquisition processing of synapsin-immunolabeled brain images obtained with a two-photon microscope. The borders of the MB lip (orange) and dense collar (green) are shown on a frontal optical section of the right median MB calyx **(A)**. The lip borders were identified in multiple optical sections of the MB calyx to reconstruct its volume **(B)**. The 40 μm thick subregion used to quantify the number of microglomeruli is highlighted in violet. Single optical sections of the right median lip are displayed before **(C)** and after **(D)** deconvolution and resampling. The upper panels represent frontal sections, with the violet line showing the position used to display the transversal sections across 40 μm in the lower panels. The squares in (**D**) represent the ROIs (10×10×10 μm^**3**^) used to compare the manual and automated quantification methods in the lip (orange) and dense collar (green). Scale bars are 100 μm **(A, B)** and 50 μm **(C, D)**.

### Consistent number of microglomeruli obtained with the automated counting in small ROIs

A major obstacle to the development of automated methods for quantifying microglomeruli was their high vicinity within the MB neuropil. Besides increasing the *z*-resolution by the application of two-photon microscopy^26^, the most important improvement in the separability of microglomeruli in the present study was obtained by applying a 3D deconvolution algorithm to the raw data with the precisely measured point spread function of the microscope objective (Supplementary Fig. S1).

The resulting resolution of the MB images was sufficient to perform an automated analysis of the number of microglomeruli in the lip and dense collar (Fig. 1C-D) using a multiple threshold segmentation.

For a given threshold of signal intensity, an automated object segmentation algorithm provided the central coordinates of each separate object in the image volume. This was repeated for various thresholds and all central coordinates were compared. The centers separated by a distance shorter than a defined optimized value were considered to belong to the same microglomerulus. The final number of distinct object centers was considered to represent the number of microglomeruli (see Materials and methods, and Supplementary Fig. S2 for a workflow). These numbers and the ones obtained by manual counting had to be corrected for the image distortion along the optical axis (Fig. 1C-D lower panels, see Materials and methods for details about the correction method) which caused additional fluorescence leaking from the outside into the ROIs. This new automated method can only be applied to images whose signal intensity does not saturate. Signal saturation can be reduced during image acquisition, and it can be avoided when selecting the ROI (see Materials and methods).

The automated and manual counting methods were applied to 1000 μm^3^ cubic ROIs located in the MB lip and dense collar (Fig. 1D). The manual method consisted in the visual identification and counting of microglomeruli, including those located on the borders of the ROI. In both MB subregions, the numbers of microglomeruli obtained with both methods were comparable (Fig. 2). A statistical analysis showed no significant difference between the two methods (paired *t-*tests; Lip: *t* = −1.34, *p* = 0.21; Dense collar: *t* = 1.25, *p* = 0.24; *n* = 10). This is the first validation of an automated method for the quantification of microglomeruli by direct comparison with the established manual method^8^.

**Figure 2.**
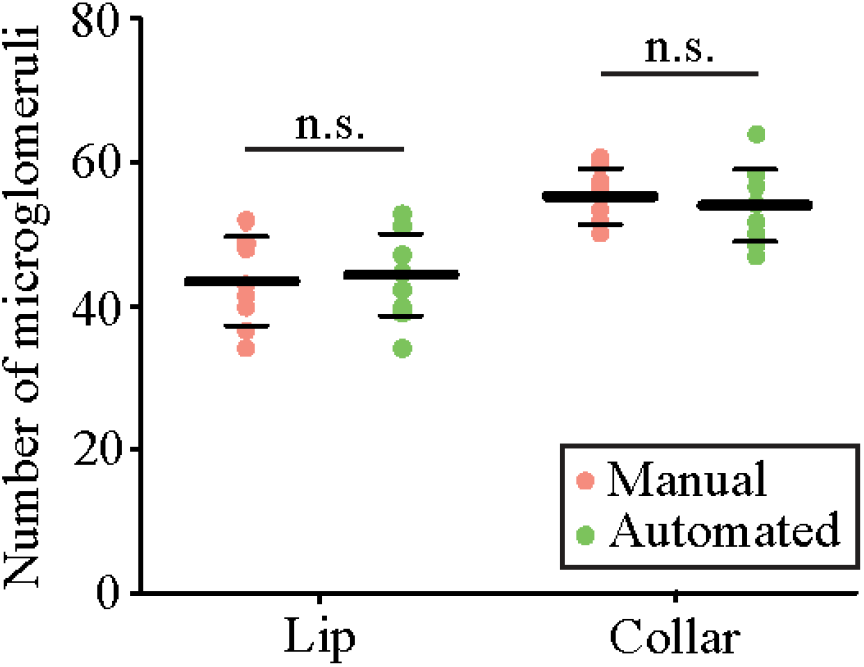
Comparison of the number of microglomeruli counted with the automated and manual methods in 10×10×10 μm^3^ ROIs positioned within the MB lip and collar of various bees. The number of microglomeruli counted with the manual method (orange) did not differ significantly from the number obtained with the automated method (green) in the lip (paired *t-*test; *t* = −1.34, *p* = 0.21) and in the dense collar (*t* = 1.25, *p* = 0.24). The thick horizontal bars represent the mean number of microglomeruli counted with each method and the thin bars show the standard deviations (*n* = 10).

Although the microglomerular organization varies greatly between insect species^28^, the present automated method is scale-free and it allows quantifying objects in any kind of system as long as the contrast is high enough to resolve neighboring objects. Depending on the vicinity and size of microglomeruli, the minimal distance used to consider segmented objects as part of the same microglomerulus needs to be adjusted (see Materials and methods).

The results obtained with the automated method in 1000 μm^3^ ROIs were also consistent with those reported in the literature by manual counting in bees of similar age (Table 1). This was not the case for previously suggested automated methods^24,27^. Peng and Yang^24^ obtained values 18-fold lower than the ones obtained by manual counting in 1000 μm^3^ ROIs (Table 1). This was partly due to the application of their method to the whole lip and not to 1000 μm^3^ ROIs, thereby considering a possible heterogeneity in the distribution of microglomeruli in this MB subregion^28^. Most of the deviation may, however, be due to imperfect segmentation methods and to the insufficient distance of 5 μm between slices used in their study^25^. Since the volume of microglomeruli ranges on average^8^ from 2.5 μm^3^ to 4 μm^3^, their coarse depth sampling may have overlooked many boutons. These shortcomings were reduced by using two-photon microscopy with finer axial sampling and by applying a 3D deconvolution to the MB images (Fig. 1), which greatly improved image resolution and was fundamental for automated discrimination of microglomeruli.

**Table 1.**
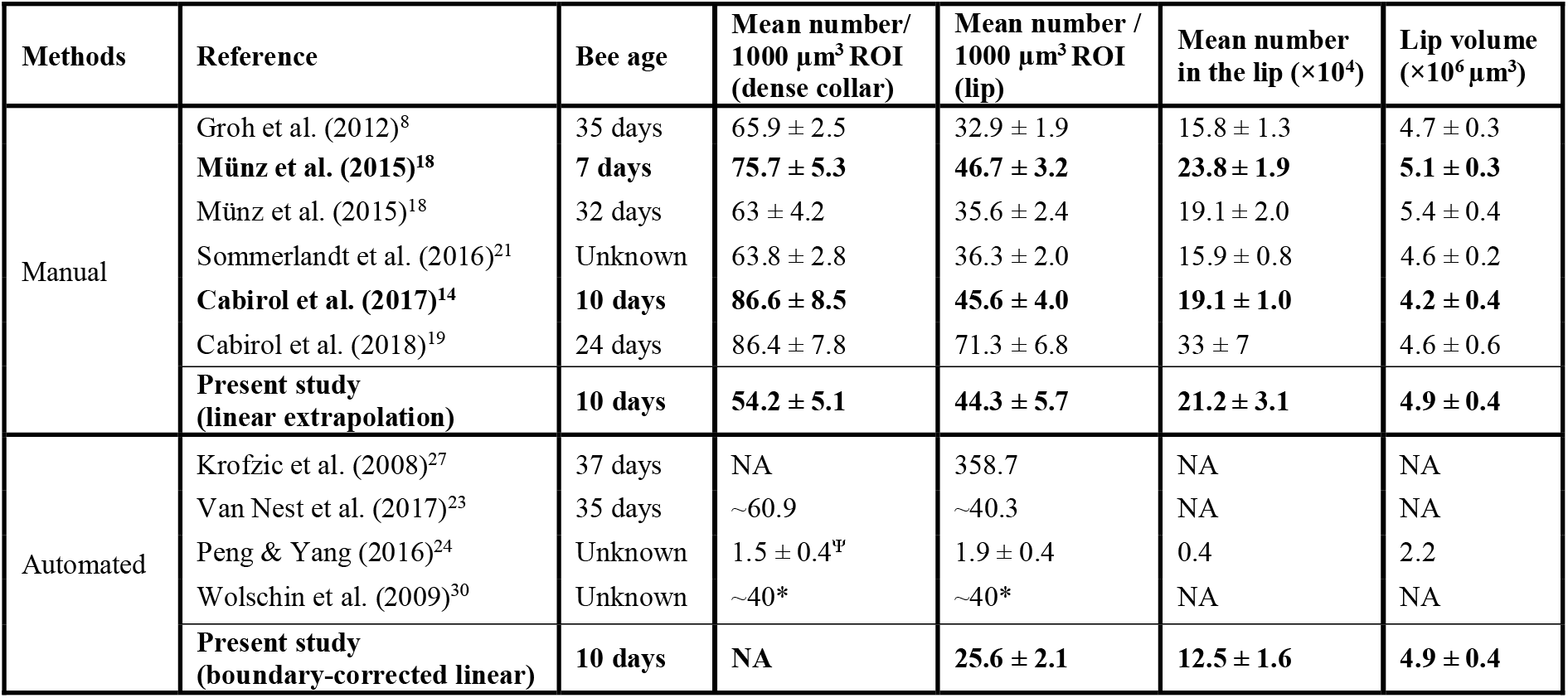
Numbers of microglomeruli reported in the literature. Manual and automated counts of synapsin-positive boutons in ROIs of 1000 μm^3^ positioned within the lip and collar are compared between studies. Values are presented as means (± standard deviation). For the manual method, the number of microglomeruli counted in the ROIs of 1000 μm^3^ was linearly extrapolated to the volume of the whole lip. The values resulting from the automated method in the present study were obtained by applying the boundary-effect model to ROIs of increasing volumes or by linearly extrapolating the number of microglomeruli counted in the 40-μm-thick lip subregion to the corresponding volumes (boundary-corrected linear). Studies performed on bees with an age similar to the present study are highlighted in bold. ^Ψ^Not only dense collar. * Lip and collar not differentiated. Adapted from Rössler *et al.* (2017)^25^.

### From ROIs to the whole lip region

Previous manual methods considered a linear relationship between the number of microglomeruli and the volume of the ROIs in which they were counted. It became a standard to extrapolate the numbers of microglomeruli counted in 1000 μm^3^ ROIs to the volume of the whole MB subregion^8^. The problem of this procedure is the inclusion of microglomeruli which intercept the ROI boundaries but have more than half their volume outside. What appears to be a negligible inaccuracy, sums up to a significant error due to the large surface-to-volume ratio in small ROIs. This error was quantified by applying the automated method to ROIs of increasing volume (Fig. 3A). The number of microglomeruli showed a drastic deviation from a linear scaling with volume. A model that assumed that the number of microglomeruli intersected by the ROI boundaries scales with the surface area of the ROIs was developed and called “boundary-effect model” (see Materials and methods). The dependence of the number of microglomeruli on the ROI volume predicted by the boundary-effect model fitted well the number obtained by automated counting in small ROIs (Fig. 3A) and in the 40-μm-thick lip subregion (Fig. 3A inset). It proves that the high accuracy of automated counting is conserved for large volumes, where a comparison with manual counting is not possible.

**Figure 3.**
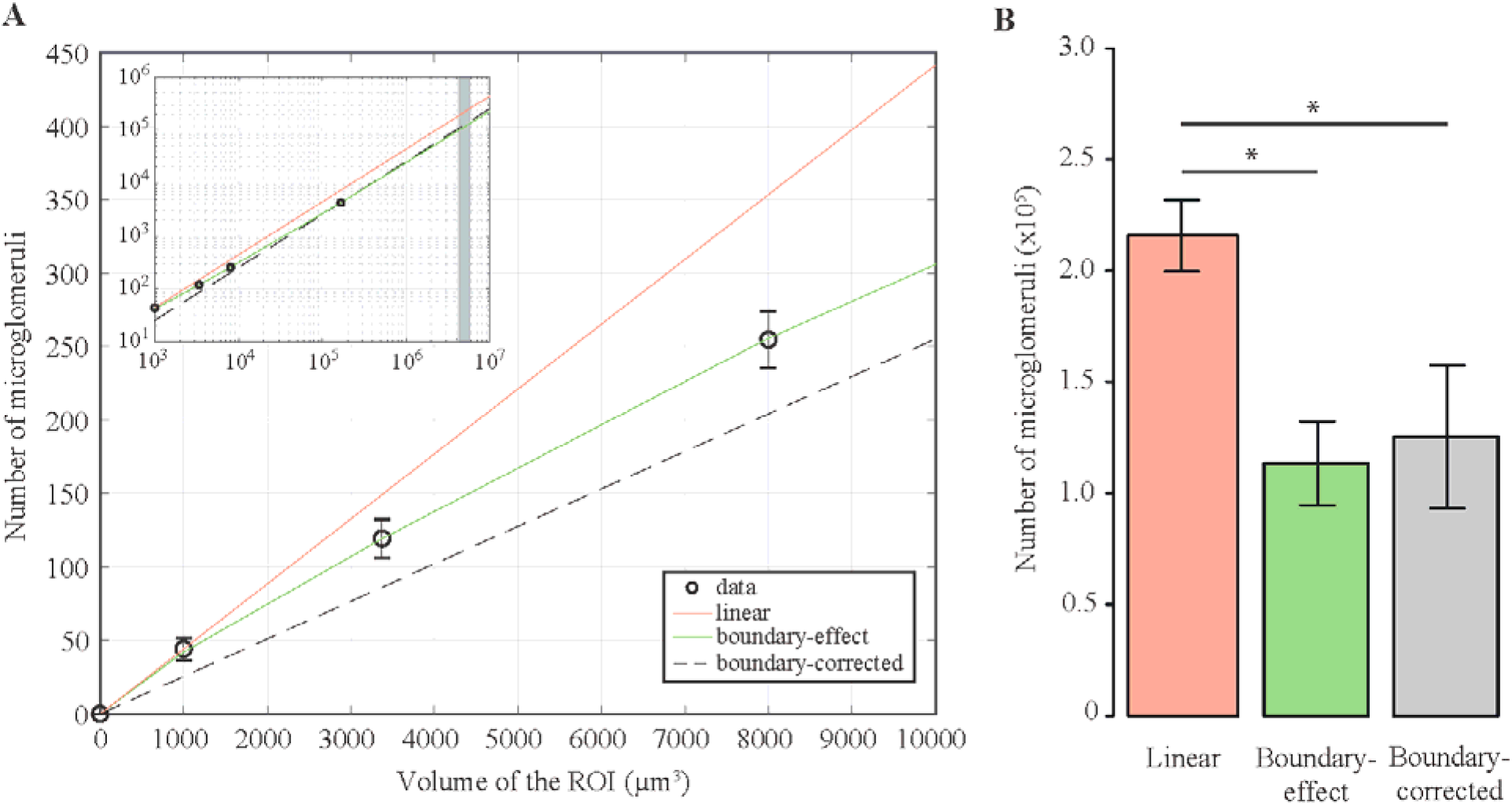
Comparison of the numbers of microglomeruli estimated in the whole MB lip using two different models. **(A)** The relationship between the number of microglomeruli counted inside a ROI and the volume of this ROI was described by fitting two models (full lines) to the experimental data obtained via automated counting (open circles). The linear model (orange), used in previous studies, consisted in a linear extrapolation of the number of microglomeruli counted in 1000 μm^3^ ROIs to the volume of the whole lip. In the boundary-effect model (green), the numbers of microglomeruli measured in four ROIs of different volumes were fitted with a function that assumes the boundary effects to scale like the ROI surface area (Formula (5) Materials and methods section). For large volumes (>10^5^ μm^3^), boundary effects became negligible and the volume dependence became linear (inset). The number of microglomeruli counted in the 40-μm-thick lip subregion, whose volume was larger than 10^5^ μm^3^, was linearly extrapolated to other volumes to obtain boundary-corrected values (dashed black line). The vertical grey bar in the inset represents the volume range of the whole lip. **(B)** The model used to define the relationship between the number of microglomeruli and the ROI volume impacted the predicted number of microglomeruli in the whole lip (repeated-measures ANOVA; *F*(1,9) = 0.81, *p* < 0.0001, *n* = 10). The linear model overestimated the number of microglomeruli in the whole lip compared to the boundary-effect model (Tukey HSD; *p* < 0.0001) and to the boundary-corrected linear extrapolation (*p* < 0.0001), which did not differ between themselves (*p* = 0.36). Error bars represent the standard deviation, \**p* < 0.0001.

Since boundary effects become negligible for volumes larger than 10^5^ μm^3^ (Fig. 3A inset), a linear extrapolation of the number of microglomeruli from the 40-μm-thick lip subregion to other volumes, hereafter called the boundary-corrected linear extrapolation, should produce reliable numbers. The number of microglomeruli in the whole lip obtained from this boundary-corrected linear extrapolation or predicted by fitting the boundary-effect model significantly differed from the number predicted by the linear model (repeated-measures ANOVA; *F* (1, 9) = 0.80, *p* < 0.0001, *n* = 10) (Fig. 3B). The linear model overestimated the number of microglomeruli on average by 70% compared to the boundary-effect model (Tukey HSD; *p* < 0.0001) and to the boundary-corrected linear extrapolation (*p* < 0.0001) (Fig. 3B). As expected, a linear extrapolation of the number of microglomeruli from the lip subregion to the whole lip produced negligible deviations from the boundary-effect model (Tukey HSD; *p* = 0.37) (Fig. 3B). This result confirms that applying the automated method to a subregion of 40 μm depth is sufficient to avoid boundary effects. Although the method was not applied to the entire dense collar, its efficiency to quantify microglomeruli in the 1000 μm^3^ ROI suggests that it would give reliable numbers in a larger ROI. As done in the lip, the ROI used to quantify microglomeruli in the dense collar should have a volume large enough to avoid boundary effects when extrapolating the resulting number to the entire structure.

A comparison between studies suggested a possible overestimation of the number of microglomeruli previously reported in the lip (Table 1). For bees with an age equivalent to the ones used in the present study (10 days), manual counting combined with the application of the linear model resulted in a considerably higher number of microglomeruli in the whole lip compared to the numbers obtained with the new automated method (Table 1, bold line below). The number of microglomeruli in the lip has been shown to decrease in older bees due to the synaptic pruning associated with foraging onset^18,19^. Yet, the number of microglomeruli reported in the whole lip of old foraging bees is higher than the one obtained in the present study in 10-day-old bees, while it is lower in 1000 μm^3^ ROIs. In one study where the manual method was applied, authors state explicitly that microglomeruli were only counted when most of their volume was falling within the ROI^8^. To avoid boundary effects this rule needs indeed to be followed with an exact threshold of 50% of the volume. Strict validation of this criterion makes manual counting even more time-consuming. However, boundary effects might therefore not be the main cause of the large discrepancy between our automated counts and previously reported manual data, and, therefore, another potential source of error is explored in the following.

### Heterogeneous spreading of microglomeruli within the lip

Advanced image processing, based on 3D deconvolution, strongly improved the quality of our images acquired with a 20× objective (Fig. 1). This allowed avoiding the use of high magnification objectives, which are not suitable for analysis of large volumes of the MB due to their small working distance and their small field of view. Thanks to the extended volume data, we were able to analyze for the first time the spatial distribution of microglomeruli in a subregion of the MB lip (Fig. 4A) and to measure their local density (Fig. 4B-D). The data revealed a substantial heterogeneity of the microglomerular distribution within the lip in all 3 dimensions.

**Figure 4.**
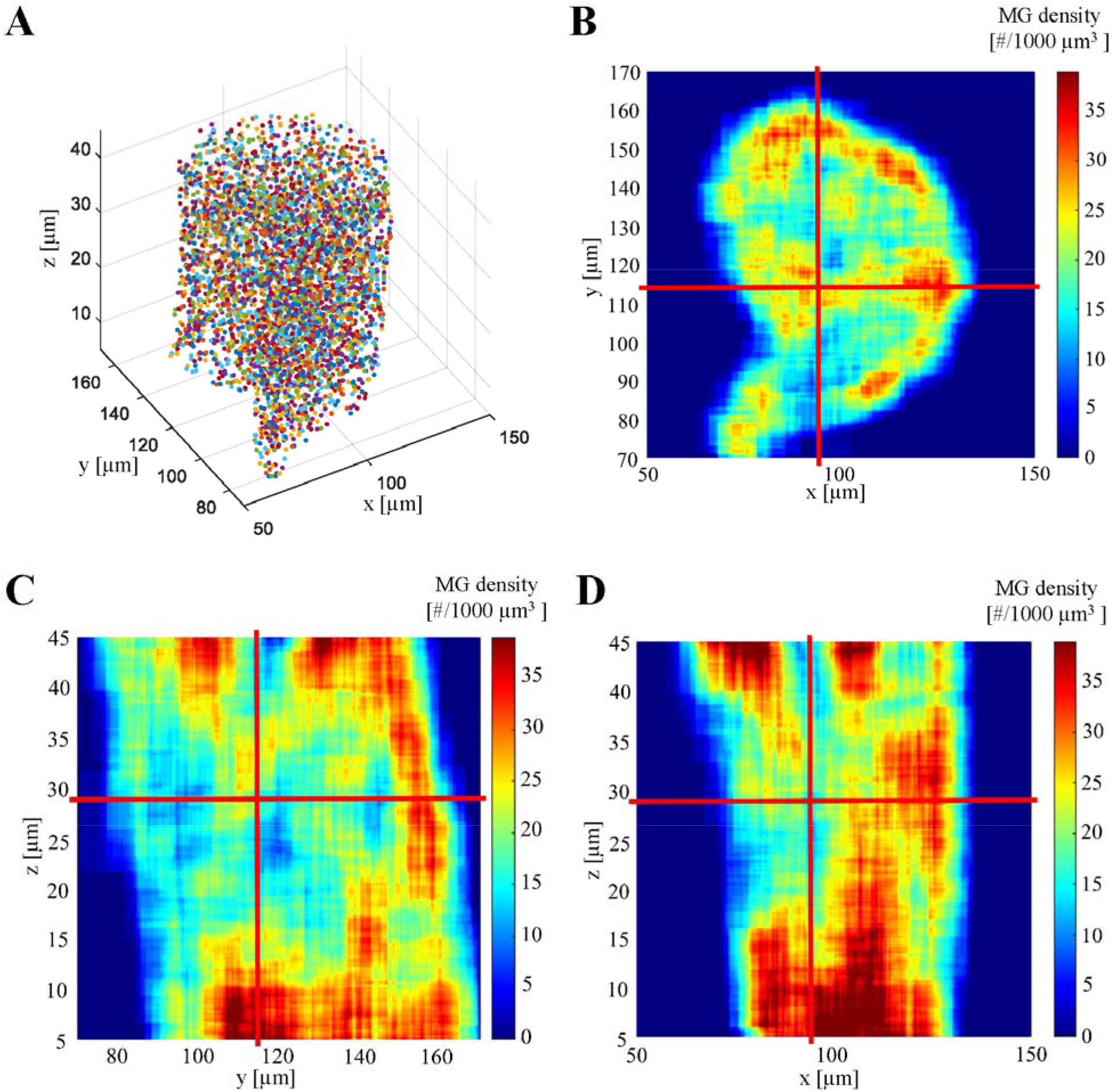
Spatial distribution of microglomeruli identified within a subregion of the right median MB lip of one individual using the automated method. **(A)** The central coordinates of microglomeruli (colored circles) were used to visualize their 3D distribution in a lip section of 40 μm thickness (random colors for visual discrimination) **(B-D)** The density of microglomeruli in #/1000 μm^3^ was obtained by a running average, using a volume element of that size. This revealed the heterogeneity in the microglomerular distribution in all dimensions of the MB lip. The red lines represent the focal planes used to display all 3 dimensions.

This exposed a further shortcoming of counting microglomeruli in ROIs, which is prone to large fluctuation depending on the ROI position. In order to estimate an average density of microglomeruli in 1000 μm^3^ ROIs, the number of microglomeruli obtained with the automated method in the 40-μm-thick lip subregion was linearly interpolated back to a 1000 μm^3^ volume. The resulting average density of 25.6 ± 2.1 per 1000 μm^3^ is not affected by the boundary effects and considers the heterogeneous microglomerular distribution. The differences between this value and densities in the MB neuropils reported in previous studies with manual counting (Table 1) might, therefore, be partially caused by the heterogeneity of the microglomerular distribution.

It should be noted, however, that part of these discrepancies between studies can be inherent to the samples. Indeed, inter-colony variation and different experimental conditions may account for differences in microglomerular numbers. Although we report a lip volume similar to previous studies, the number of microglomeruli within lips of identical volume can still be different because the volume of the MB calyx is also influenced by the size of the dendritic tree formed by MB intrinsic neurons^29^.

Given the heterogeneous spreading of microglomeruli within the lip, the major advance of our method compared to previous automated methods^24,27,30^ is its success in extracting accurately the number of microglomeruli and their 3D coordinates from images obtained with a 20× objective, whose working distance and field of view allow to image larger volumes. A current limit to the application of this automated method to the entire lip is the computational power required for the multiple threshold segmentation and the identification of microglomeruli. Future improvements to the method should, therefore, focus on overcoming this limit. Yet, the number of microglomeruli estimated in the entire lip with the automated method showed little variation between bees despite differences in the location of the 40-μm-thick lip subregion (see standard deviations in Table 1). This suggests that the number of microglomeruli counted in a 40-μm-thick lip subregion might be similar across the lip.

When smaller structures need to be studied (*e.g.* individual synapses), higher resolution and higher magnifications are required. Until now, this has mainly been achieved using electron microscopy^8,31–33^. However, new advances in light microscopy, in particular, the development of nanometer-resolution microscopes, will allow for future optical studies down to the synaptic level^34^. Still, the described problem of boundary effects, being a general phenomenon, will be relevant also for those applications. Even in completely different scenarios, whenever objects of an extended size are counted within small sample volumes, a linear extrapolation of these counts should be examined critically for potential biases from boundary effects, as reported in this study.

## Conclusions

The present study provides a new automated method for synaptic bouton quantification in larger brain regions than what could previously be achieved. The application of the method to such brain volumes avoided one bias that occurs when synaptic boutons are counted in smaller regions of interest: the boundary effects, which are due to boutons located on the ROI’s borders. It also revealed the heterogeneous distribution of synaptic boutons within the brain region. Future studies should aim at extending the imaging volume to be able to analyze the spatial structure of synaptic boutons in the entire mushroom body, to investigate *e.g.* the stereotypy of these patterns, their symmetry^35^, and their plasticity.

## Materials and methods

### Animals

Experiments were performed in June 2018 on a honey bee colony (*Apis mellifera*) maintained at the University of Trento in Rovereto. Inter-individual variability in brain structure was reduced by using same-age honey bees reared under controlled conditions. For this, a comb of brood about to emerge was taken from the hive and left in complete darkness in an incubator (34°C, 55% humidity) overnight. Newborn adult bees were collected in the following morning. They were placed in cages (8×5×4.5 cm, 15 bees per cage) in complete darkness in an incubator (34°C, 55% humidity) for 7 days with unlimited access to sucrose solution (50% w/w) and water. The sucrose solution and water were changed every two days.

### Immunostaining of synapsin in whole-mount brains

The protocol used for the immunostainings is fully described elsewhere^8^. The brains of 7-day-old honey bees (*n* = 10) were dissected in Ringer solution (130 mM NaCl, 5 mM KCl, 4 mM MgCl_2_, 5 mM CaCl_2_, 15 mM Hepes, 25 mM glucose, 160 mM sucrose; pH 7.2) and fixed overnight in 4% formaldehyde at 4°C on a shaker. Brains were rinsed with PBS (1 M; 3×10 min) and permeabilized successively in 2% PBS-Triton X100 (PBS-Tx; 10 min) and 0.2% PBS-Tx (2×10 min). Brains were blocked in 2% Normal Goat Serum (in 0.2% PBS-Tx) for 1 h at room temperature. They were then incubated with the α-synapsin antibody SYNORF1 for 3 days at 4°C on a shaker, rinsed with PBS (5×10 min), and incubated with the Goat anti-Mouse secondary antibody Alexa-488 conjugate for 3 days at 4°C on a shaker. Finally, the brains were rinsed with PBS (5×10 min) and dehydrated in an ascending ethanol series (30, 50, 70, 90, 95, 100, 100, 100%; 10 min each). Clearing and mounting were performed in methyl salicylate.

### Image acquisition

Whole-mounted brains were imaged using a two-photon microscope (Ultima IV, Bruker) with excitation at λ = 780 nm (Ti:Sa laser, Mai Tai Deep See HP, Newport), using a 20× objective (NA 1.0, water immersion, Olympus)^36^. For volume measurements of the MB lip, image stacks of the entire right median calyx were acquired with a resolution of 512×512 pixels with a pixel size of 1.03×1.03 μm. The inter-slice interval was 5 μm. For the quantification of microglomeruli, a subregion of the outer lip of the right median calyx was imaged with a pixel size of 0.34×0.34 μm, over a depth of 50 μm and with an inter-slice interval of 0.5 μm. The intensity of the laser was changed with depth to compensate for the scattering loss which increases exponentially with imaging depth *z*. Therefore, at the image stack’s upper limit *z*_0_ a laser power *P*_0_ was chosen to produce an unsaturated image of optimal contrast. At the final imaging depth *z*_*f*_, the laser power *P*_*f*_ was adapted to produce the same overall fluorescence intensity, again without saturating pixels. The laser power was then automatically adapted following the equation:

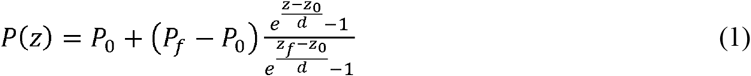

The parameter *d* defines a characteristic compensation length, which has been optimized so that the fluorescence remains constant across depth^37^. Signal saturation was thereby avoided across the entire stack, allowing the automated method to perform an artifact-fee segmentation of the microglomeruli. If an optimal compensation cannot be achieved, the laser power might be set at intermediate layers and interpolated according to the above formula. A link to the raw data can be found in the Supplementary information (Supplementary Material S3).

### Volume reconstruction of the MB lip

The volume measurements of the lip were semi-automatically performed using AMIRA (V5.4, Thermo Scientific). Using the SegmentationEditor, lip borders were manually defined every 5-6 slices and interpolated by the software. After carefully checking the accuracy of the delimitation on each slice, the volume of the lip was calculated using the MaterialStatistics function. The metadata was exported in XML-format for further statistical analyses.

### Image processing for the quantification of microglomeruli

To improve contrast and resolution of the image stacks, essential for robust discrimination of microglomeruli, images of the lip subregion were post-processed by a 3D deconvolution in AMIRA. The required Point Spread Function (PSF) of the microscope objective was measured using fluorescent beads (TetraSpeck, 0.1 μm, Thermo Fisher) (Supplementary Fig. S1). The maximum-likelihood deconvolution algorithms required 100 iterations for sufficient convergence. Images were then resampled to a final voxel size of 0.1×0.1×0.1 μm (Fig. 1).

### Quantification of microglomeruli in ROIs

The same deconvoluted and resampled images of the lip were used to compare the numbers of microglomeruli obtained with the manual and automated methods. The two methods were first applied to ROIs of size 10×10×10 μm positioned within the lip (Fig. 1D).

Manual counting was performed using the AMIRA LandmarkEditor by placing landmarks on visually identified microglomeruli (Fig. 5D).

**Figure 5.**
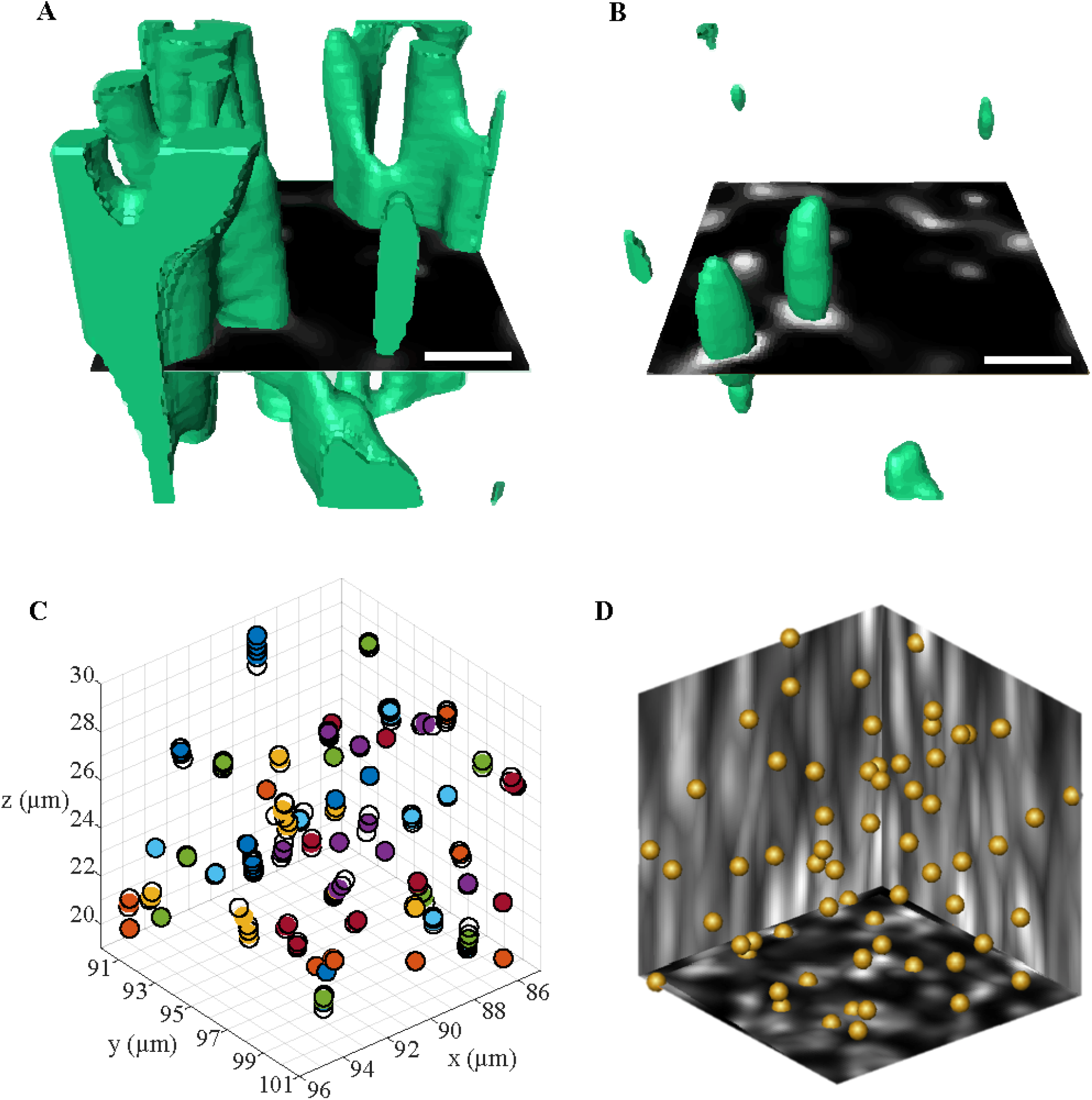
Quantification of microglomeruli in a ROI (10×10×10 μm^3^) positioned within the MB lip. In the automated method, a multiple threshold segmentation was applied to label voxels having an intensity level above a defined value (from 20% to 90% of the maximum intensity in steps of 10%). Two examples show a single optical section and the 3D objects formed by the connected voxels segmented with the thresholds of 20% **(A)** and 90% **(B)** (green labels). **(C)** The central coordinates of these 3D objects (open circles) were compared, and those with centers distant less than 0.8 μm were considered representations of the same microglomerulus (colored circles). **(D)** In the manual method, landmarks were placed on top of visually identified microglomeruli and automatically counted by the software. Scale bars are 2 μm.

The automated counting protocol was based on the idea that repeated image segmentation using varying thresholds of signal intensity assures that all separate objects are extracted from the image at least once. Objects selected by different thresholds and having a similar location were considered to belong to the same microglomerulus. In details, the AMIRA SegmentationEditor was used to label voxels whose signal intensity was above a certain threshold. The threshold was varied in repeated applications from 20% to 90% of the maximum image intensity in steps of 10% (Fig. 5A, B). For each threshold, connected labeled voxels formed objects whose center coordinates and volumes were extracted using the RegionStatistics module and exported in XML-format for further analyses.

Via a code written in MATLAB (R2018, MathWorks) (Supplementary Methods S4), objects larger than 10 μm^3^ were removed from all threshold datasets because they most likely contained multiple microglomeruli. A minimum size limit of 0.05 μm^3^ was applied to the lowest threshold dataset only (20% of signal maximum) to remove artefacts. Since connected objects separate and have reduced sizes when increasing the threshold, they should not be considered as artefacts for higher thresholds. The identification of microglomeruli was performed by calculating the pair-wise 3D distance between all object centers. Objects whose centers were below a distance of 0.8 μm were considered to represent the same microglomerulus. This parameter, as well as the minimum and maximum object’s volume, were optimized regarding the robustness with which the algorithm reproduced manual counting results for all samples. First, microglomerular numbers obtained by varying these parameters were compared to the ones counted beforehand with the manual method. Then, a visual comparison (Fig. 5C, D) confirmed the efficiency of the automated method to identify microglomeruli.

The same automated method was then applied to images of the lip subregion. To remove ringing artifacts due to the deconvolution, 5 μm had to be cropped at the *z*-boundaries, resulting in a lip subregion of 40 μm depth. The volume of this lip subregion was calculated as described for the whole lip.

### Correction for the elliptic image of the microglomeruli

The number of microglomeruli obtained in all ROIs and in the lip subregion had to be corrected because our imaging setup distorted the images of microglomeruli along the optical axis labeled here as *z*. This was due to a refractive-index mismatch between the water-immersion objective and the mounting media of our samples. Image segmentation using MATLAB was used to determine the correct microglomerular size of 0.86 ± 0.53 μm (mean ± std) along the transversal dimensions and 3.3 ± 1.3 μm along the optical axis. This means that signals from microglomeruli located in a layer of approximately *r*_*leak*_ = 1,16 ± 0.89 μm above and below the ROIs along the optical axis were leaking into the ROIs and biased the counts. Absolute sizes must be taken with care when comparing them to microglomeruli images obtained by other methods because the dehydration procedure strongly shrinks the tissue^10,35^ and the deconvolution algorithm reduces the resolution-limited object sizes.

Given the size of the layer causing the bias, we were able to obtain a corrected number of microglomeruli *N*_*corr*_ from the raw numbers *N*_*raw*_ starting from the following formula:

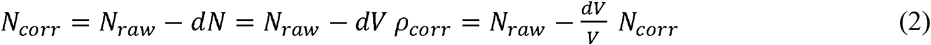

where *dN* is the number of microglomeruli located above or below the ROI whose signal is leaking inside, which can be quantified by the relative affected volume 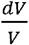 in the ROI, *V* being the ROI volume. 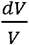 equals the ratio of leaking depth *r*_*leak*_ = 1.16 μm to the size of the ROI *d*_*ROI*_ multiplied by two because upper and lower surface both contribute, which gives the final correction formula:

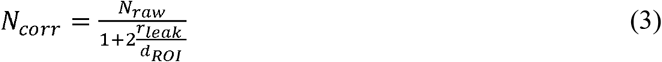

This reduced the measured numbers of microglomeruli by 19% in the 1000 μm^3^ ROIs, 13% in the 3375 μm^3^ ROIs, 10% in the 8000 μm^3^ ROIs, and 5% in the lip subregion of 40 μm depth. These corrections were applied to all manually and automatically obtained counts.

### Quantification of boundary effects when counting in small ROIs

Even in non-distorted images, some microglomeruli are inevitably located at the boundaries of the ROIs. A correct density value should consider only the microglomeruli that have at least half their volume inside the ROI. Instead, both our automatic counting and the established manual counting protocols consider all microglomeruli intersecting the ROI boundaries as inside. To quantify the overestimation of the microglomerular density due to this effect, one starts in the same way as above by estimating the affected volume (2). But now the effect occurs not only on the upper and lower boundary but on all 6 surfaces of a cubic ROI. The affected depth from the surface equals to the radius of the imaged microglomeruli *r*_*MG*_ so that instead of (3) we obtain the correction formula:

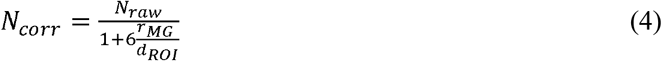

To test this model, ROIs of different volumes: 1000 μm^3^ (10×10×10 μm^**3**^), 3375 μm^3^ (15×15×15 μm^**3**^), and 8000 μm^3^ (20×20×20 μm^**3**^) were selected around the same center point. The included microglomeruli were counted with the automated method.

The numbers of microglomeruli were averaged over different bees and their scaling as a function of ROI volume was analyzed by fitting two curves. First, a linear model assumed that boundary effects were negligible by extrapolating the density *ρ* of microglomeruli (the number measured in a 1000 μm^3^ ROI) linearly with the volume *V*:

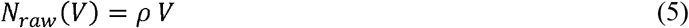

Second, we used the boundary-effect model (4) and expressed the correct number of microglomeruli in terms of density and volume and the ROI size by the cubic root of its volume: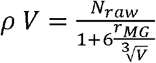. Solving for *N*_*raw*_, we obtain the fitting function:

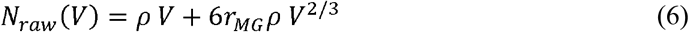

which shows that besides the correct number of microglomeruli which increases linearly with the ROI volume, the number of erroneously counted microglomeruli located at the ROI boundaries scales as *V*^2/3^. This boundary effect loses importance with increasing *V*, showing a linear scaling for high *V*. Comparing the real data to the two models allowed to evaluate whether the data actually deviated from the linear behavior and if so, to identify the ROI volume from which boundary effects became negligible.

### Statistical analyses

The normality of data distribution and the homogeneity of variances were confirmed for all variables by applying a *Shapiro-Wilk test* (*p* > 0.05) and a *Bartlett test* (*p* > 0.05) respectively. A paired *t*-test was used to compare the number of microglomeruli counted with the manual and automated methods in the ROIs of 1000 μm^3^. The numbers of microglomeruli estimated in the whole lip by the three methods were compared by applying an ANOVA for repeated measures, followed by a Tukey HSD post-hoc test.

## Data availability

The datasets generated and/or analysed during the current study are available from the Image Data Resource: https://idr.openmicroscopy.org/ or from the corresponding author on request.

## Additional information

**Supplementary information** accompanies this paper.

## Competing interests

The authors declare no competing interests.

## Author contributions

AC and AH designed research; AC performed research and analyzed data; AC and AH wrote the paper.

## Acknowledgments

This work was funded by the Autonomous Province of Bolzano (Project B26J16000310003) https://r1.unitn.it/stefanie/. The authors thank the members of the Neurophysics group at the Center for Mind/Brain Sciences & Department of Physics for useful remarks on manuscript. They also thank an anonymous reviewer for important comments and suggestions on the paper.

