## Supplementary information for "Automated quantification of synaptic boutons reveals their 3D distribution in the honey bee mushroom body"

**Supplementary Figure S1. The two-photon imaging system's point spread function.** Images of 0.1 µm fluorescent beads are displayed along the (x,y)-axis **(A)** and (y,z)-axis **(B)**. An average over several beads was used to measure the Point Spread Function of the microscope. The resulting measures were the radial Gaussian width $\sigma_{xy} = 0.39 \pm 0.01$ µm and the axial Gaussian width $\sigma_z = 2.3 \pm 0.1$ µm. The voxel size is 0.13×0.13×0.5 $\mu m^3$.

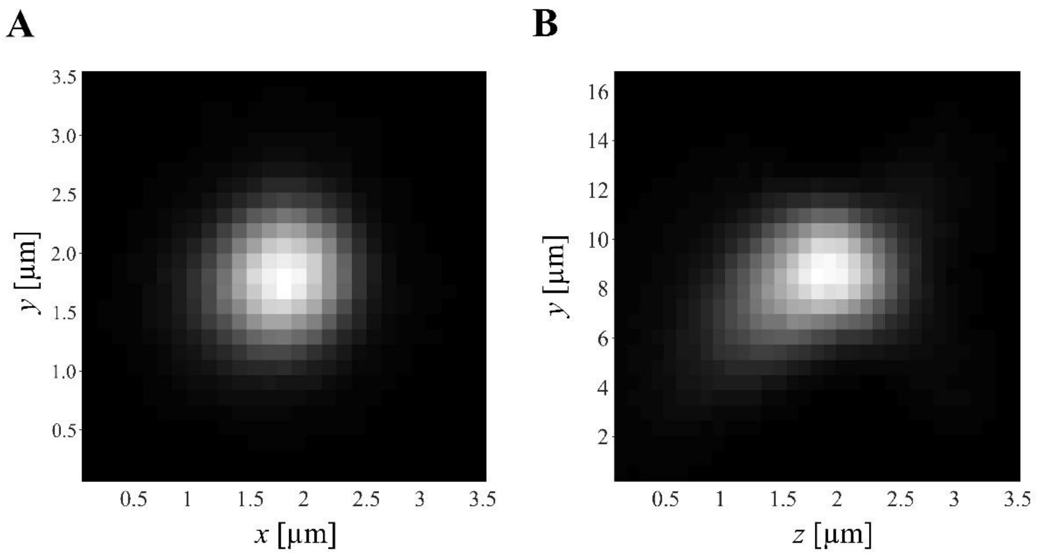


**Supplementary Figure S2. Workflow of the successive processes used to extract the number and 3D distribution of microglomeruli in the mushroom body lip from images acquired with a two-photon microscope.** Two sets of images of the lip were obtained with different acquisition parameters optimizing the volumetric measurements of the whole region (pink) or the automated identification of microglomeruli in a subregion of the lip (green). Some processes were applied to both sets of images (orange). The method required the use of AMIRA (V5.4) to extract data from the images and of MATLAB (R2018) to further analyze these data. The number of microglomeruli counted in the lip subregion, whose volume has also been measured, can finally be linearly extrapolated to the volume of the entire lip.

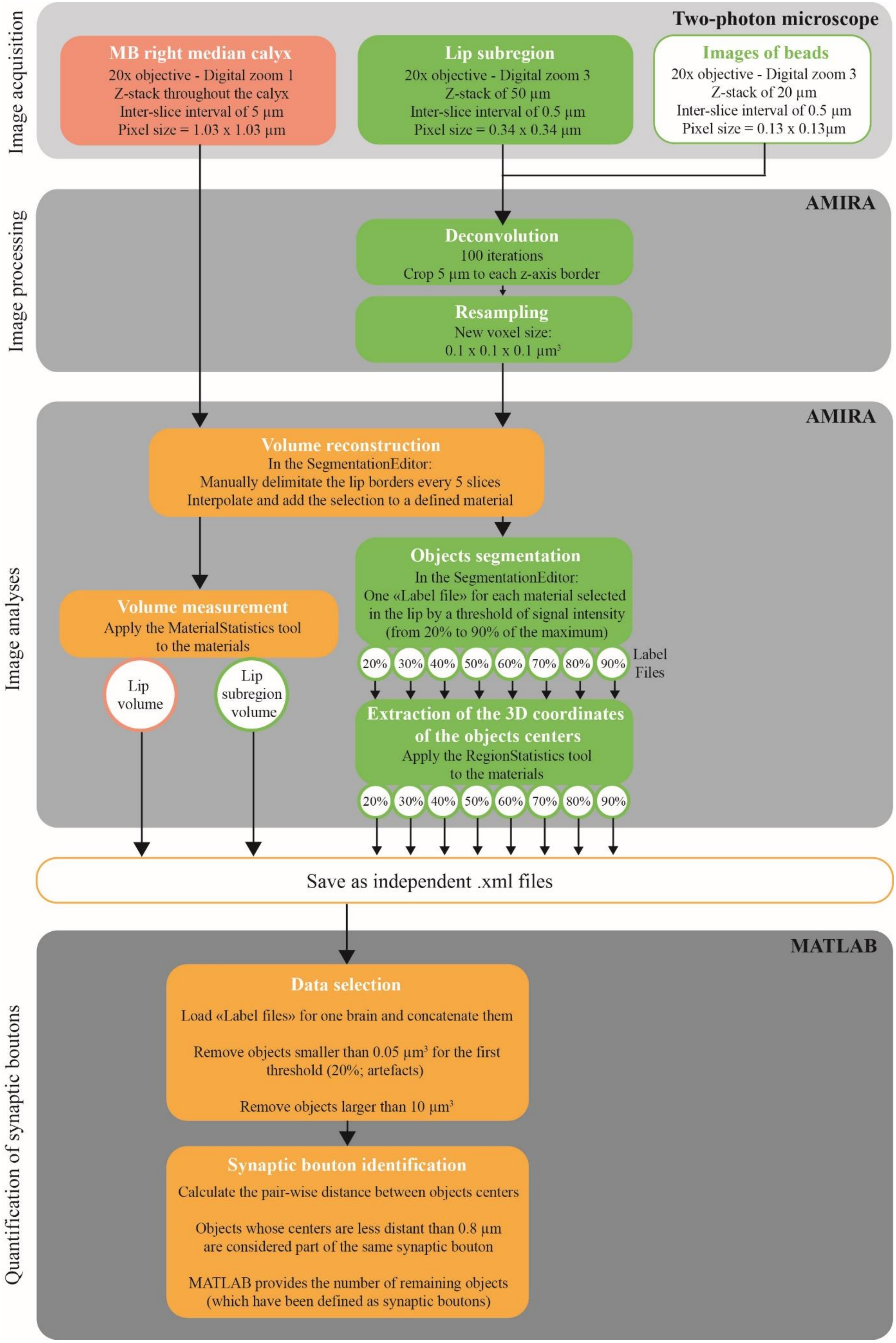

Image acquisition
Two-photon microscope
MB right median calyx
20x objective - Digital zoom 1
Z-stack throughout the calyx
Inter-slice interval of 5 µm
Pixel size = 1.03 x 1.03 µm
Lip subregion
20x objective - Digital zoom 3
Z-stack of 50 µm
Inter-slice interval of 0.5 µm
Pixel size = 0.34 x 0.34 µm
Images of beads
20x objective - Digital zoom 3
Z-stack of 20 µm
Inter-slice interval of 0.5 µm
Pixel size = 0.13 x 0.13µm
Image processing
AMIRA
Deconvolution
100 iterations
Crop 5 µm to each z-axis border
Resampling
New voxel size:
0.1 x 0.1 x 0.1 µm³
Image analyses
AMIRA
Volume reconstruction
In the SegmentationEditor:
Manually delimitate the lip borders every 5 slices
Interpolate and add the selection to a defined material
Objects segmentation
In the SegmentationEditor:
One «Label file» for each material selected
in the lip by a threshold of signal intensity
(from 20% to 90% of the maximum)
Volume measurement
Apply the MaterialStatistics tool
to the materials
20% 30% 40% 50% 60% 70% 80% 90%
Label Files
Lip volume
Lip subregion volume
Extraction of the 3D coordinates of the objects centers
Apply the RegionStatistics tool
to the materials
20% 30% 40% 50% 60% 70% 80% 90%
Save as independent .xml files
Quantification of synaptic boutons
MATLAB
Data selection
Load «Label files» for one brain and concatenate them
Remove objects smaller than 0.05 µm³ for the first threshold (20%; artefacts)
Remove objects larger than 10 µm³
Synaptic bouton identification
Calculate the pair-wise distance between objects centers
Objects whose centers are less distant than 0.8 µm are considered part of the same synaptic bouton
MATLAB provides the number of remaining objects (which have been defined as synaptic boutons)

**Supplementary Material S3. Image-stacks of 40 µm-thick lip subregions from ten honey bee brains.** Whole-mounted brains were imaged using a two-photon microscope after immunostaining synapsin. Images were acquired with a 20 × objective (NA 1.0, water immersion, Olympus), a pixel size of 0.34×0.34 µm, and an inter-slice interval of 0.5 µm.
All raw data image files and the corresponding microglomeruli coordinate maps will be available from the Image Data Resource: https://idr.openmicroscopy.org/

**Supplementary Methods S4. Matlab script that identifies individual objects via their coordinates obtained by multiple thresholding.** This script requires as input .xml files containing the 3D coordinates of objects, obtained by image segmentation at different intensity threshold levels via image analysis software *e.g.* AMIRA. It removes too large objects from all threshold analyses and too small objects from the lowest threshold analysis. It performs a pairwise comparison of the object coordinates to identify as distinct objects those with coordinates separated by more than a minimum distance. It visualizes their spatial distribution and their density profile in all 3 dimensions.

```
%% Creation of the matrix containing all objects coordinates.
% Load the files containing the 3D coordinates of all objects
clear all; close all
path='\\Path-leading-to-the-folder-containing-the-xml-files\';
files=dir([path '*.xml']);
% Concatenate all threshold files
Araw=[]; Region=[];
for j=1:size(files,1)
    [num,txt,raw]=xlsread([path files(j).name]);
    Araw=cat(1,Araw,num(:,4:7));
    Region=cat(1,Region,txt(2:end,2));
end

%% Define the parameters used to identify microglomeruli.
distlim=0.8; % Distance between the centers of two objects under which they
are considered as being the same microglomerulus
volmin=0.05; % Minimum volume applied to the first threshold of signal
intensity
volmax=10; % Maximum volume applied to objects selected by all thresholds
of signal intensity
lim=[50 150 70 170 5 45]; % Minimal and maximal limits of the x-, y- and z-
axes

%% Remove objects that are not microglomeruli but either the neuropil
region
%  in which they are located, or the area surrounding the neuropil
respectively
%  named "Lip" and "Ext" in this example.
A=Araw;
for i=1:size(Araw,1)
    if Region{i}(1:3)=='Ext'|Region{i}(1:3)=='Lip'
        A(i,:)=nan;
        Region{i}=nan;
    else A(i,:)=Araw(i,:);
    end
end
A(any(isnan(A),2),:)=[];
Region(any(cellfun(@(Region)any(isnan(Region)),Region),2),:)=[];

%% Volume thresholds
% Set a minimum volume for the 20% threshold only(removing artifacts)
A2=[];
for i=1:size(A,1)
    if (Region{i}(1:4)=='MG20')&(A(i,1)<volmin)
        A2(i,:)=nan;
```

```
    else A2(i,:)=A(i,:);
    end
end
A2(any(isnan(A2),2),:)=[];
% Set a maximum volume for all thresholds (removing highly connected
% microglomeruli)
A2(any(A2(:,1)>volmax,2),:) = [];
A2=A2(:,2:4); % Working copy without the volume of objects

%% Identify objects selected by multiple thresholds using the pairwise
distance
%  between their centers and consider them as being the same
microglomerulus
A3=A2; % Working copy
ED = pdist(A2); % Pairwise distance
sqED = squareform(ED); % In matrix form
same = sqED < distlim; % distance smaller than distlimit
overl= sum(same,2); % Column vector of number of overlaps
m=0; % Cluster count
while ~isempty(overl)
    m=m+1; % Cluster count
    [val,id]=sort(same(1,:),'descend'); % id positions of overlap, val is 1
for overlap
    B(m,:)=mean(A3(id(1:overl(1)),1:3),1); % Average position of overlaps
    pop(m)=overl(1);
    A3(id(1:overl(1)),:)=[]; % Remove treated cluster
    same(id(1:overl(1)),:)=[];
    same(:,id(1:overl(1)))=[];
    overl= sum(same,2); % Recalculate remaining overlaps
end;

%% Projected density: Estimate the number of microglomeruli in a 1000ùm3
%  cubic ROI centered around each voxel.
bins=[1000 1000 400];
x = linspace(lim(1),lim(2),bins(1));
y = linspace(lim(3),lim(4),bins(2));
z = linspace(lim(5),lim(6),bins(3));
ed2Dx = {y,z};
ed2Dy = {x,z};
ed2Dz = {x,y};

% Calculate 2D density pattern in 10 microns depth along the x-axis
B2=B;
for i=1:size(B,1)
    if (B(i,1)<100)&&(B(i,1)>90) % Define the minimum and maximum x-
coordinates between which density will be calculated
        B2(i,:)=B(i,:);
    else B2(i,:)=nan;
    end
end
B2(any(isnan(B2),2),:)=[];
x2 = hist3([B2(:,2),B2(:,3)],'edges',ed2Dx);
x3 = movmean(x2,100,1,'omitnan');
x5 = movmean(x3,100,2,'omitnan');
x5=x5.*10000;

% Calculate 2D density pattern in 10 microns depth along the y-axis
B3=B;
for i=1:size(B,1)
    if (B(i,2)<120)&&(B(i,2)>110) % Define the minimum and maximum y-
coordinates between which density will be calculated
```

```
        B3(i,:)=B(i,:);
    else B3(i,:)=nan;
    end
end
B3(any(isnan(B3),2),:)=[];
y2 = hist3([B3(:,1),B3(:,3)],'edges',ed2Dy);
y3 = movmean(y2,100,1,'omitnan');
y5 = movmean(y3,100,2,'omitnan');
y5=y5.*10000;

% Calculate 2D density pattern in 10 microns depth along the z-axis
B4=B;
for i=1:size(B,1)
    if (B(i,3)<35)&&(B(i,3)>25) % Define the minimum and maximum z-
coordinates between which density will be calculated
        B4(i,:)=B(i,:);
    else B4(i,:)=nan;
    end
end
B4(any(isnan(B4),2),:)=[];
z2 = hist3([B4(:,1),B4(:,2)],'edges',ed2Dz);
z3 = movmean(z2,100,1,'omitnan');
z5 = movmean(z3,100,2,'omitnan');
z5=z5.*10000;

%% Graphic representation
% plot 3D: Represents the centres of microglomeruli within the lip
subregion
figure,subplot(2,2,1),hold on
scatter3(A2(:,2),A2(:,1),A2(:,3),25,'k') % Original MG
colormap(lines(size(B,1)))
scatter3(B(:,1),B(:,2),B(:,3),10,(1:size(B,1))','filled') % Clustered MG
hold off
grid on, axis(lim),axis square,view([142.5,45])
xlabel('x [{\mu}m]');ylabel('y [{\mu}m]');zlabel('z [{\mu}m]');
title(['3D MG Centers: ' num2str(m) ' clusters'])

% 2D densities of microglomeruli in the lip subregion and in all 3
dimensions
subplot(2,2,2)
pcolor(x,y,z5')
colormap jet, colorbar, shading flat,axis square, axis(lim(1:4))
xlabel('x [{\mu}m]');ylabel('y [{\mu}m]');
title('MG density')
cl = caxis

subplot(2,2,3)
pcolor(y,z,x5')
colormap jet, colorbar,shading flat, axis square
xlabel('y [{\mu}m]');ylabel('z [{\mu}m]');
title('MG density')
axis(lim(3:6))
caxis(cl)

subplot(2,2,4)
pcolor(x,z,y5'),colormap jet, colorbar,shading flat,axis square
xlabel('x [{\mu}m]');ylabel('z [{\mu}m]');
title('MG density')
axis([lim(1:2) lim(5:6)])
caxis(cl)
```
